# *In Vivo* Multiplexed Modeling Reveals Diverse Roles of the TBX2 Subfamily and *Egr1* in *Ras*-Driven Lung Adenocarcinoma

**DOI:** 10.1101/2025.03.15.642187

**Authors:** Athar Khalil, Trang Dinh, Meaghan Parks, Rebecca C. Obeng, Berkley Gryder, Adam Kresak, Yuxiang Wang, Jeff Maltas, Madeline Bedrock, Xiangzhen Wei, Zachary Faber, Mira Rahm, Jacob Scott, Thomas LaFramboise, Zhenghe Wang, Christopher McFarland

**Affiliations:** Department of Genetics and Genome Sciences, Case Western Reserve University School of Medicine, Cleveland, OH 44106, USA; Cancer Genomics and Epigenomics Program, Case Comprehensive Cancer Center, Case Western Reserve University, Cleveland, OH 44106; Department of Pathology, Case Western Reserve University School of Medicine, Cleveland, OH 44106, USA; Department of Pathology, University Hospitals Cleveland Medical Center, Cleveland, OH 44106, USA; Translational Hematology & Oncology Research. Cleveland Clinic. Cleveland, OH 44106. USA

## Abstract

The TBX2 subfamily of T-box transcription factors (including *Tbx2*, *Tbx3*, *Tbx4*, *Tbx5*) plays an essential role in lung development. Downregulation of these genes in human Lung adenocarcinoma (LUAD) suggests that these genes may be tumor suppressive, however because downregulation appears to occur primarily via epigenetic change, it remains unclear if these changes causally drive tumor progression or are merely the consequence of upstream events. Herein, we developed the first multiplexed mouse model to study the impact of TBX2 subfamily loss, alongside associated signaling genes *Egr1*, *Chd2*, *Tnfaip3a*, and *Atf3*, in *Ras*-driven lung cancer. Using TuBa-seq, a high-throughput tumor-barcoding system, we quantified the growth effects of these knockouts during early and late tumorigenesis. *Chd2* loss consistently suppressed tumor progression, while *Tbx2* loss exhibited stage-dependent effects. Notably, *Egr1* emerged as a potent tumor suppressor, with its knockout increasing tumor size (∼5x) at 20 weeks, surpassing *Rb1* loss. Transcriptomic analyses of *Egr1*-deficient tumors suggested immune dysregulation, including heightened inflammation and potential markers of T cell exhaustion in the tumor microenvironment. These findings indicate that *Egr1* may play a role in suppressing tumor growth through modulating immune dynamics, offering new insights into the interplay between tumor progression and immune regulation in LUAD.

## Introduction

Lung adenocarcinoma (LUAD) is a genetically diverse cancer driven by numerous oncogenic and tumor-suppressive events(1). The T-box gene family, comprising eighteen members in mammals, encodes transcription factors that are crucial for embryonic development and tissue homeostasis(2,3). The four members of the TBX2 subfamily (*Tbx2*, *Tbx3*, *Tbx4*, and *Tbx5*) play an indispensable role in lung development, with their expression being crucial for various aspects of normal lung morphogenesis and differentiation(4–7). In previous studies, we observed marked downregulation of the TBX2 subfamily genes in clinical LUAD specimens. The downregulation of these genes appears to be predominantly mediated by epigenetic mechanisms, particularly DNA methylation dysregulation(8,9). Because these clinical analyses are inherently correlative, it remains unclear whether the downregulation of these genes causally drives tumor progression or is merely a consequence of broader transcriptional reprogramming. Reactivation of these genes in LUAD cell lines led to reduced cell growth, suggesting a potential tumor suppressive function(10). This effect is amplified in the presence of a Kirsten rat sarcoma viral oncogene homolog (*KRAS)* mutation– the most common oncogenic event in human LUADs(11). In other cancer types, TBX2 genes play a differential role in regulating cell cycle progression, proliferation, senescence, apoptosis, inflammation and metastasis(12). With such a diverse and central role in cancer, these genes were shown to function as either oncogenes or tumor suppressors (TSs) depending on the cellular context (Fig1A)(13–15). Hence, faithful models of TBX2 gene loss in an *in vivo* lung environment are needed to understand the causal role of these genes in LUAD, particularly in the context of *Kras*-driven oncogenesis.

Herein, we developed the first multiplexed mouse model to study TBX2 subfamily loss along with potential downstream effector genes (*Egr1, Chd2, Tnfaip3a, and Atf3)* identified through our *in vitro* studies in *Ras*-driven lung cancer(10). Our approach combines <u>tu</u>mor-<u>ba</u>rcoding with high-throughput barcode <u>seq</u>uencing (Tuba-seq)(16). Tumors initiation was carried out using pools of barcoded Lenti-sg*RNA*/Cre viruses, and the resulting tumors were analyzed at two distinct time points thereby comprehensively assessing multiple stages of tumorigenesis, including tumor initiation, growth, and progression of Kras^G12D^ driven lung tumors, across nine distinct genotypes. Notably, early growth response 1 (*Egr1*) emerged as a critical regulator, with its loss having a more pronounced impact on tumor progression than the loss of the well-established tumor suppressor gene Retinoblastoma 1 (*Rb1*).

## Results

### 1 Development of the first *in vivo* lung cancer model of TBX2 subfamily signaling associated genes using multiplexed CRISPRko and lineage-tracing

To enable CRISPR/Cas9-mediated somatic genome editing within the context of oncogenic *Kras* variant, we crossed mice with Cre/lox-activated alleles of KRAS ^G12D^ (Kras^LSL-G12D/+^) and Cre/lox-activated Cas9 (R26^LSL-Cas9-eGFP/+^), hereafter named a KC mouse model(16,17). Tumors were initiated through intratracheal intubation with barcoded pooled-Lenti-sgRNA/Cre vectors targeting the four TBX2 genes (*Tbx2*, *Tbx3*, *Tbx4*, and *Tbx5*) and four of their associated effector genes identified previously through our *in vitro* transcriptome profiling (*Egr1*, *Chd2*, *Tnfaip3a*, and *Atf3*) (Fig. 1A and B)(11). The utilized lenti-sgRNA-Pool/Cre pool included control genome-targeting inerts (sg*Neo1* and sg*Neo2*), positive and negative controls targeting the known tumor suppressor gene *Rb1* and the essential gene *Pcna*, respectively. Each gene was targeted with two unique Lenti-sgRNA/Cre vectors optimized for on-target cutting specificity within the first three exons of the targeted gene (Supplementary Table S1). To investigate the role of these genes at both early and late tumorigenesis stages, cohorts of mice were evaluated at 6- and 20-weeks post-intubation, respectively (Fig. 1B). Lung weights and histological examination revealed fewer visible tumors at 6-week, despite these mice receiving higher viral titers (Fig. 1C and D, Supplementary Fig. S1A). Lung tumors observed at both time points included hyperplastic lesions and adenomas with foci of high-grade/malignant features, all of which stained positively for the lung lineage-defining transcription factor NKX2.1/TTF-1(Fig. 1C).

**Figure 1:**
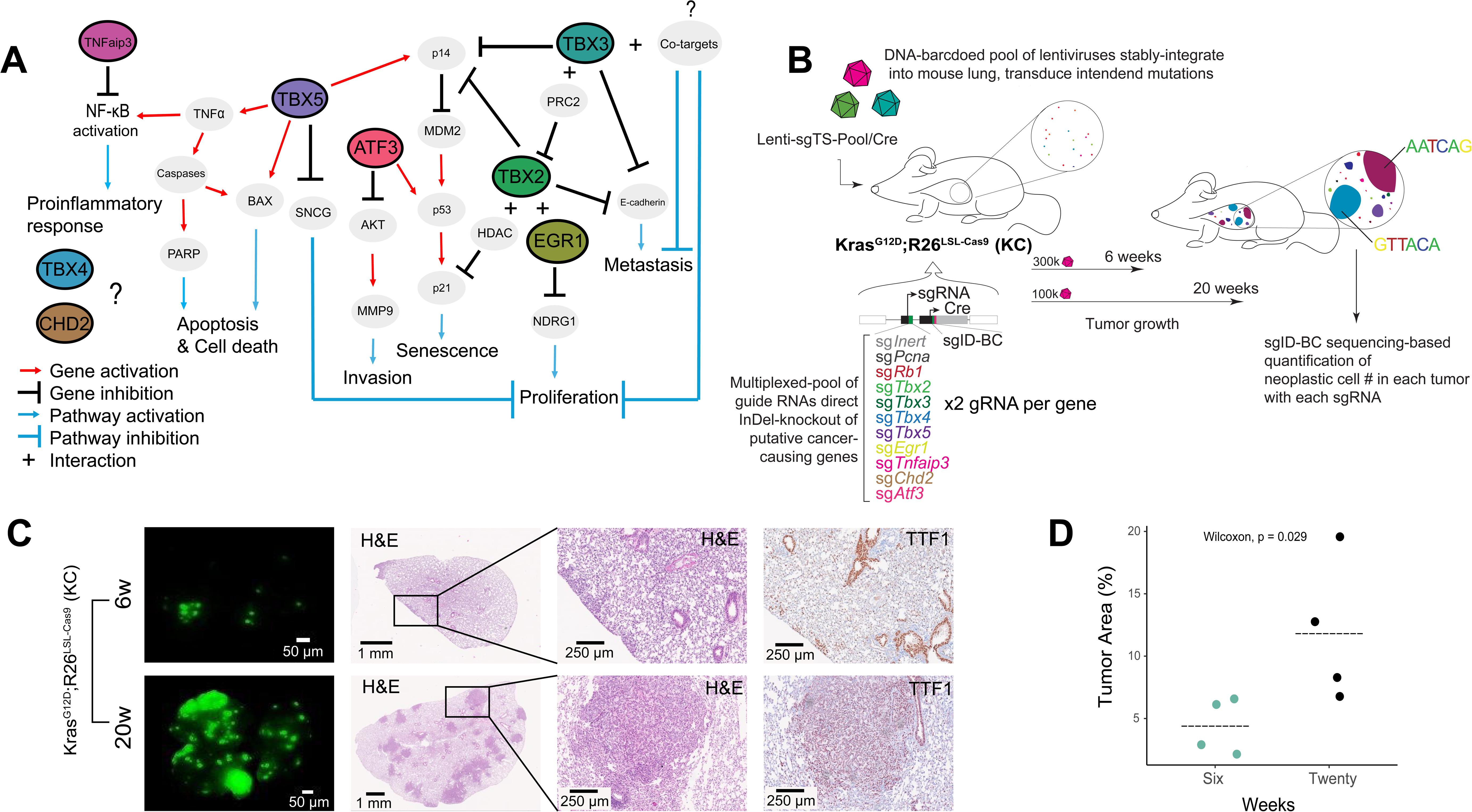
TBX2 Signaling in cancer and Tuba-seq approach for evaluating their role in *Kras*-driven LUADs. A) Schematic representation of the molecular basis of TBX2-associated signaling pathways in cancer. *TBX2*, *TBX3*, and *TBX5* contribute to the dysregulation of multiple cancer hallmarks, including cell proliferation, senescence, apoptosis, invasion, metastasis, and the pro-inflammatory response. *EGR1*, *TNFaip3*, *ATF3*, *Tbx4, and Chd2* are all modulated by TBX2 signaling in lung cancer cell lines and recurrently dysregulated in human lung cancers. While *EGR1*, *TNFAIP3, and ATF3* all directly modulate key carcinogenic pathways, the mechanisms by which *TBX4* and *CHD2* influence cancer remain largely unexplored(15,22,40–49). *AKT* (AKT serine/threonine kinase), *BAX* (BCL2 associated X, apoptosis regulator), *HDAC* (Histone deacetylase), *MDM2* (MDM2 proto-oncogene, E3 ubiquitin protein ligase), *MMP9* (Matrix metallopeptidase 9), *NDRG1* (N-myc downstream-regulated gene 1), *p14ARF* (Cyclin-dependent kinase inhibitor 2A, isoform p14ARF), p*21CIP1* (Cyclin-dependent kinase inhibitor 1A), *p53* (Tumor protein p53), *PAsrp* (Phosphatidylserine receptor protein), *PRC2* (Polycomb repressive complex 2), *SNCG* (Synuclein gamma), and *TNFa* (Tumor necrosis factor alpha). B) Experimental schematic of the Tuba-seq approach used to investigate combinatorial inactivation of potential LUAD regulators in vivo. Tumor initiation was achieved through intratracheal intubation with Lenti-sgTS-Pool/Cre in Kras^LSL-G12D^,Rosa26^CAG-LSL-Cas9-GFP^ mice. The Lenti-sgTS-Pool contained two inert sgRNA vectors along with a positive control (sg*Rb1*) and a negative control (sg*Pcna*). Each of the eight genes studied was targeted using two distinct sgRNAs. Each sgRNA vector included a unique sgID and a random barcode, allowing quantification of individual tumor sizes via deep sequencing. The genotype, time points and lentiviral titers are as indicated. C) Representative images of lung lobes from mice at 6- and 20-weeks post-tumor initiation. Images include fluorescence dissecting scope views, H&E-stained sections, and TTF1-stained (lung epithelial marker) sections of lung lobes. Scale bars are indicated on each image. D) Quantification of the percent tumor area in representative mice revealed a significant increase at 20 weeks compared to 6 weeks (Wilcoxon test, p<0.05). Each dot represents an individual mouse, with horizontal bars indicating the mean tumor area.

### 2 Multiplexed quantification of TBX2 genes function in *Kras*-driven lung tumors

To precisely quantify the growth effects of each gene knockout within the same mouse at all potential tumor sizes – from tens of cells to millions, we stably labeled the genetically diverse tumors with lentiviral-mediated DNA barcodes (termed Tumor Barcoding, TuBa-seq). Tumor sizes and quantities were determined *en masse* by analyzing genomic DNA extracted from bulk tumor-bearing lung tissue, followed by deep sequencing of the double barcode region, which identifies both the short guide RNA (sgID) and descendants (‘million’ BarCode, mBC) of every transduced cell. Sequencing performed to an average depth of >10^7^ reads per mouse allowed us to reliably quantify tumor sizes below 400 cells. To translate barcode read tallies to absolute tumor sizes, we spike-in DNA barcodes of known cell number prior to cell lysis. After barcodes are tallied, we then eliminate spurious tumors and potential technical replicates by generating a statistical model of sequencing errors and barcode diversity (Supplementary Fig. S1B, C and D). We assessed the impact of knocking out each of the potential tumor suppressor genes (TSG) by analyzing the distribution of absolute cell numbers for each gene knockout (Fig. 2A). Consistent with histological evaluations, the mean total cell number initiated by either Lenti-sgInerts or Lenti-sgTSGs was significantly higher in the 20-week cohort compared to the 6-week cohort (Supplementary Fig. S1E). The average total cell number produced by a single sgInert was significantly smaller than the combined mean cell number produced by our studied knockouts, indicating an overall marked acceleration in cellular proliferation.

**Figure 2:**
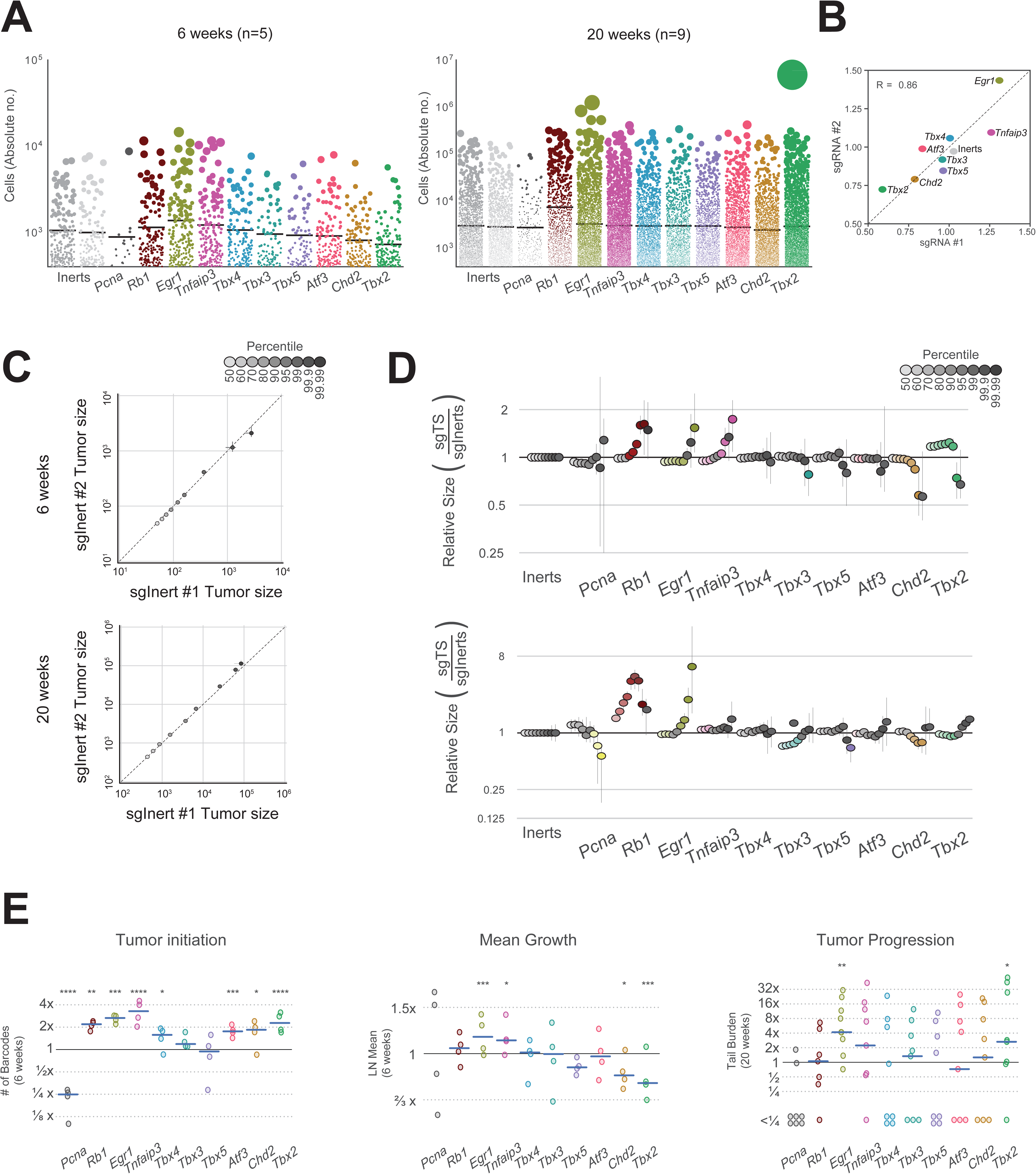
Multiplexed somatic CRISPR–Cas9 genome editing reveals that loss of *Egr1* has the most pronounced and consistent effect on tumor initiation, growth, and progression in *Kras*-driven lung cancer. A) Individual tumor sizes ordered by mean effect of targeted gene knockout in KC mice at 6- and 20-weeks post tumor initiation. Number of mice in each cohort is as indicated. Each dot represents a tumor. The area of each dot is proportional to the number of cancer cells in each tumor. B) Mean effect of gene knockout of each sgRNA in KC mice at 6-weeks post tumor initiation. LN Mean is a Maximum Likelihood Estimator (MLE) of mean tumor size under a Log-Normal sampling distribution. Each gene in this study was targeted by two high-specificity sgRNAs imparting consistent growth effects (Pearson’s *R* = 0.86, P = 0.0029, Methods). C) Quantile-Quantile (Q-Q) plot of tumor size distributions of the top 50^th^ – 99.99^th^ percentiles of the two Inert sgRNAs at both 6 and 20 weeks demonstrates reproducibility of size profiles. D) Analysis of relative tumor sizes in KRAS driven tumors 6 and 20 weeks post tumor initiation. Relative size of tumors for each targeted gene (sgTS) is divided by the respective sg*Inerts* percentile and merged across replicate mice. Percentiles significantly different from sg*Inerts* are in color (*P* < 0.05, two-sided bootstrap resampling). The darker the shade of color the larger the percentile, as shown in the legend in gray scale. Error bars denote 95% confidence intervals also determined by bootstrap sampling. E) Summary of three distinct growth phenotype profile. For each genotype, the number of tumors observed at six weeks (Tumor Initiation), alongside MLE of mean size at six weeks (Tumor growth, see c), and the size of the largest tumors at 20 weeks (advanced progression) is depicted. All statistics are divided by their respective values for sg*Inerts*. *P* < 0.05, 0.01, 0.001, 0.0001 for *, **, ***, and **** respectively (two-sided bootstrap resampling).

Lineage tracing uncovered a wide spectrum of effects on tumorigenesis and variability in tumor sizes with a unique profile of growth effects for each TSG knockout (Fig. 2A). Both the mean estimate and 95th percentile of tumor sizes of each gene knockout was highly correlated between the two sgRNAs targeting a gene (r = 0.87 for mean estimate, Fig. 2B, Supplementary Fig. S1F). Strong correlation of the percentile spectrums of the two Inert sgRNAs for replicates at both 6 and 20 weeks demonstrates excellent reproducibility of our growth profiles (Fig. 2C). Similarly, A quantile-quantile (Q-Q) comparison of each gene knockout against the wild-type (*Kras*-only) tumor size distribution in each mouse summarizes growth effects across all tumor sizes (Fig. D, Supplementary Fig. S2A). As cataloged previously via TuBa-seq (17), a gene’s effects on tumorigenesis reproducibly alters (i) the number of tumor lineages transduced by the same lentiviral pool, (ii) the mean/median size of a transduced tumor, and (iii) the probability of exceptionally large tumors. These exceptionally large tumors cannot be explained by a single-step Markov process, nor mode of gene-editing (Floxed-allele or CRISPRko) and are consistent with (epi)genetic progression over the time course (16). Hence, we quantified tumor progression via three distinct, previously-vetted progression measures: (i) Tumor Initiation effect (number of barcodes observed), (ii) Mean Growth effect (a Maximum Likelihood Estimator of the mean based on a Log-normal distribution, LN Mean, Supplementary Fig. S2B), and (iii) effect on Advanced Progression (total burden of tumors larger than sizes expected from a Log-normal distribution, Methods). Summary statistics for each gene were reported relative to wild-type summary statistics within each mouse, thereby controlling for extensive variability in tumor growth observed between mice (Fig. 2E) (16).

Tumor necrosis factor alpha-induced protein 3 (*Tnfaip3*)-deficient cells showed a marked increase in tumor size after 6 weeks of growth, suggesting a role in early tumor growth but not necessarily in advanced progression. (P < 0.05 for 95, 99th percentiles, and LN Mean, bootstrap resampling, Methods) (Fig. 2D). Tnfaip3, also known as A20, is a key anti-inflammatory enzyme and a critical regulator of inflammation homeostasis. Prior studies have demonstrated that the intrinsic loss of A20 in tumor cells markedly enhances lung tumorigenesis and impairs CD8+ T cell-mediated immune surveillance in both patients and murine model(18).

The profile of *Tbx2*-deficient cells exhibited a unique pattern, with an increase in tumor initiation capacity and relative sizes of smaller percentiles, accompanied by a significant *decrease* in overall LN mean at 6 weeks and the largest percentiles (Fig. 2D). Furthermore, this knockout led to the formation of some of the largest tumors observed in our screen at 20 weeks. Countervailing context-dependent effects of gene knockouts on growth have been observed previously and are consistent with known *Tbx2* biology. Inappropriate activation of *Tbx2* is thought to contribute to tumor progression by overriding senescence, thereby sustaining tumor growth. *TBX2* overexpression has been observed in pancreatic, colorectal, melanoma, endometrial, ovarian, and cervical cancers, where it contributes to tumorigenesis(15). Conversely, other lung and skin cancer models showed that *Tbx2* overexpression inhibits cell growth while being associated with increased resistance of tumor cells to the anti-cancer drug cisplatin– consistent with a highly contextual role of *Tbx2* in cancer progression(19).

Loss of *Tbx3*, *Tbx4*, *Tbx5* and *Atf3* exhibited only marginal growth effects in Ras-driven tumors, similar to their marginal effects on absolute cell number (Fig. 2D and E). *Chd2* loss consistently resulted in suppressed cellular proliferation with a reduction in total cell number at top percentiles and a significant decrease in LN mean at the studied time point. In general, inactivation of CHD family members has been implicated in various human cancers. Notably, CHD2 has been suggested to play a role in preventing breast cancer initiation. In lung cancer, *CHD2* mRNA expression has been linked to cancer stage, particularly in lung squamous cell carcinoma(20).

Lastly, deactivation of *Egr1*, a *Tbx2* partner and a direct regulator of key tumor suppressors such as *Tgfβ1*, *Pten*, and *Tp53*, increased mean growth rates at both early and late time points and resulted in exceptionally large tumors at 20 weeks post-initiation (∼5x size increase, p<0.05), surpassing even the effect of *Rb1* knockout(21). Strikingly, only *Egr1* loss significantly promoted all three growth phenotypes, exhibiting this screen’s strongest advanced progression phenotype (Fig. 2D&E, Supplementary Fig. S2B).

### 3 Immune Pathway Modulation and Tumor Progression Following *Egr1* Suppression in *Kras*-Driven Lung Cancer

To validate the tumor-suppressive role of *Egr1* and investigate its mechanism of action in a *Kras*-driven background, we generated *Egr1*-deficient tumors in KC mice by delivering Lenti-sg*Egr1*-mBC/Cre pools and compared them to *Egr1*-intact control tumors induced in KC mice using Lenti-sgInert-mBC/Cre pools (Fig. 3A). Twenty weeks after tumor initiation, *Egr1*-deficient mouse tumors developed larger tumors, compared to *Egr1*-control (Fig. 3B, Supplementary Fig. S3A).

**Figure 3:**
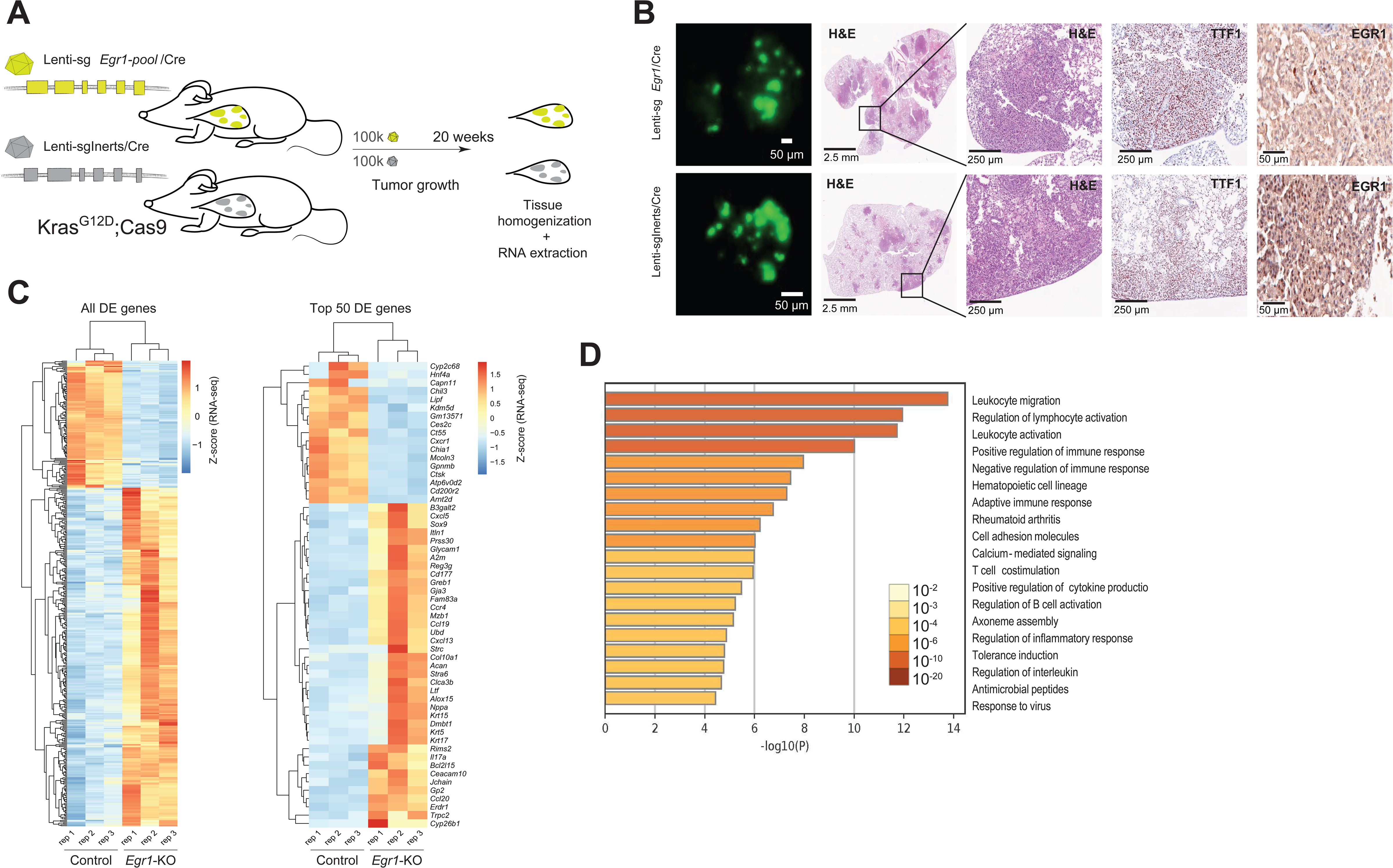
Loss of *Egr1* Promotes Tumor Progression, Immune Modulation, and Differential Gene Expression in *Kras*-Driven Lung Cancer. A) Schematic illustration of the experimental design for *Egr1* knockout versus control. KC mice were administered a pool of two sgRNAs targeting *Egr1* (n=3) or a pool of control sg*Inerts* (n=3). (B) Representative images of lung lobes from mice 20 weeks post-tumor initiation via Lenti-sg*Egr1*-pool/Cre or Lenti-sg*Inerts*. Images include fluorescence views under a dissecting microscope, H&E-stained sections, TTF1-stained sections to confirm lung adenomatous origin, and Egr1 staining to confirm knockout efficiency. C) Bulk RNA-seq analysis was conducted on lung tissues 20 weeks post-tumor initiation in *Egr1* knockout and control mice. The left panel displays a heatmap of 421 differentially expressed genes (DEGs) identified with an FDR q < 0.05, illustrating two distinct expression clusters. The right panel highlights a heatmap of the top 50 most deregulated genes, showing clear differences between the two groups. D) Gene Ontology analysis was performed using Metascape on significantly upregulated genes between *Egr1* knockout and control mice. Statistically enriched terms were identified based on cumulative hypergeometric p-values and enrichment scores. The top 20 most significant terms, predominantly associated with immune processes, are visualized.

To investigate the molecular mechanisms of *Egr1*-driven tumor growth, whole transcriptome comparison between tumor-bearing *Egr1*-knockout lungs and control mice was performed (Supplementary Fig. S3B). 421 genes were significantly deregulated in *Egr1*-knockout lung tissues as compared to control samples. Unsupervised hierarchical clustering of gene expression profiles revealed distinct differential expression patterns between the groups (Fig. 3C). Nearly all the top upregulated genes were closely linked to immune responses, inflammation, and chemokine signaling, including *Il17a*, *Itln1*, *Cd177*, *Gp2*, *Cxcl5*, and *Cxcl13*. Gene Ontology (GO) pathway analysis of the differentially upregulated genes corroborated the immune-modulatory effects of *Egr1* loss, with leukocyte migration and activation emerging as the most enriched ontology clusters (Fig. 3D). Building on these findings, Maximal Clique Centrality (MCC) analysis further pinpointed the top 12 hub genes within the most enriched immune-related pathways. These genes played critical roles in immune regulation and pathway interconnectivity across the eight primary immune ontology categories (Fig. 4A). Furthermore, Gene Set Enrichment Analysis (GSEA) revealed significant enrichment in three immune and inflammatory signaling hallmarks: allograft rejection, interferon gamma response, and interferon alpha response, collectively suggesting that *Egr1* deficiency leads to a more immunogenic tumor phenotype. Conversely, although less pronounced, several hallmark tumorigenesis pathways, including Myc, mTOR, and ROS signaling, were downregulated in these tumors (Supplementary Fig. S3C).

**Figure 4:**
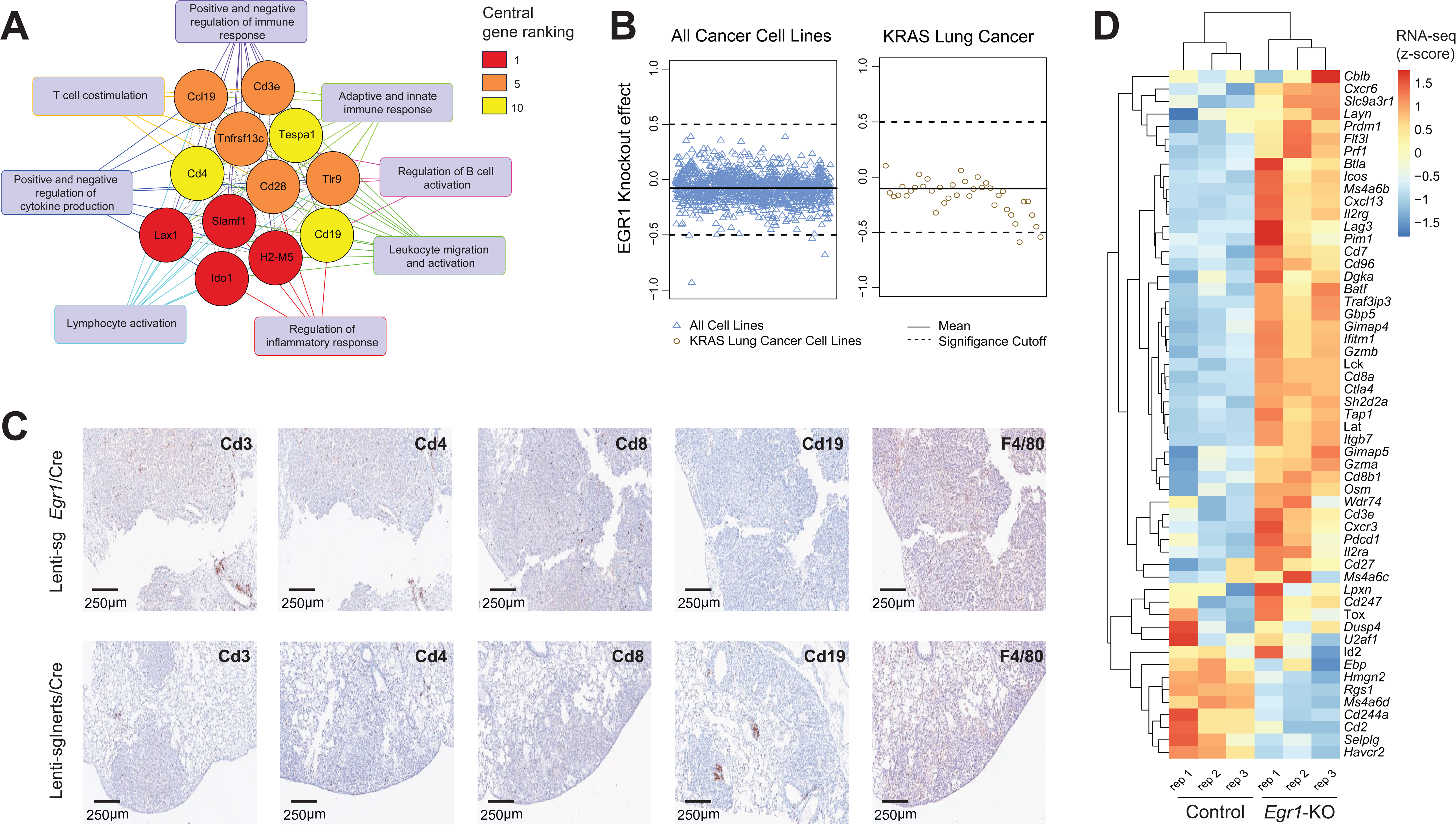
Impact of *Egr1* knockout on *in vitro* lung cancer proliferation, *in vivo* T cell infiltration, and T cell exhaustion in the tumor microenvironment. A) The immune network and top 12 hub genes are visualized. From the top 30 most significantly upregulated pathway clusters, 18 immunity-related terms and their associated genes were selected for Maximal Clique Centrality analysis using the CytoHubba plugin in Cytoscape. Gene interactions within the network indicate their participation in shared pathways. The background color of each gene box represents its centrality ranking: four genes co-ranked 1st (red), five genes co-ranked 5th (orange), and three genes co-ranked 10th (yellow). The 16 pathways associated with these 12 hub genes were further consolidated into 8 primary categories, represented by blue rectangular boxes. (B) Analysis of the DepMap CRISPR–Cas9 cancer dataset evaluating the effect of *Egr1* knockout in 342 human cancer cell lines, including independent assessments in KRAS-driven lung cancer lines, showed no significant impact on cell proliferation in either case. (C) Immunohistochemical staining of Cd4, Cd8, Cd3, Cd19, and F4/80 in *Egr1*-deficient lung tumors compared to control tumors. Scale bars are provided for each image. (D) A heatmap illustrating the expression patterns of 55 curated T-cell exhaustion markers (CellMarker 2.0 and 10X Genomics) in control and *Egr1*-deficient tumors. Differential expression analysis reveals that most exhaustion markers are upregulated in *Egr1*-deficient samples. The color scale represents relative expression levels, ranging from red (high expression) to blue (low expression).

We employed three complementary approaches to further explore the impact of *Egr1* knockout on lung tumorigenesis and the role of the tumor microenvironment in this process. First, using the Pan-Cancer Analysis of Whole Genomes (PCAWG) dataset, we demonstrated significant suppression of *EGR1* expression in clinical adenocarcinoma tissues compared to normal lung tissues (Supplementary Fig. S3D). Next, we quantified the growth effects of *EGR1* knockout in human KRAS-driven LUAD cell lines and observed no significant growth differences in the absence of an immune microenvironment (Fig. 4B, Supplementary Table S2). Additionally, immunohistochemical (IHC) staining of lung tissues revealed a marked increase in T cell infiltration in *Egr1*-KO tumor tissues, with robust infiltration of both helper T cells and cytotoxic T cells, indicating an immune response linked to *Egr1* loss (Fig. 4C). In contrast, macrophage staining intensity was similar between control and knockout samples, and B cell staining, marked by CD19, was comparable in both groups, although more aggregates were observed in the control samples (Fig. 4C).

To further explore the molecular signature of T cell status in the tumor microenvironment of *Egr1*-deficient samples, we focused on gene sets associated with T cell exhaustion. *Egr1* knockout samples exhibited substantial differential expression, including upregulation of key T cell exhaustion markers such as *Pdcd1* (PD-1), *Cd244* (2B4), *Lag3*, and *Ctla4* (Fig. 4E). These findings suggest a potential transcriptional shift suggestive of T cell exhaustion in the absence of *Egr1*. However, it is important to note that further detailed analysis using single-cell RNA sequencing or spatial transcriptomics would provide a more comprehensive understanding of the T cell status within the tumor microenvironment, which we plan to address in future studies.

## Discussion

Most putative driver mutations are altered in <10% of tumors, highlighting the need to understand the functional importance of additional actionable drivers that may be infrequently altered at the genetic level, but often deregulated through expression changes(17). Furthermore, the impact of co-occurring tumor suppressor gene alterations is often overlooked in the molecular classification of tumors. Our study addresses the limitations of current methodologies, which predominantly focus on single-driver oncogenic mutations, thereby neglecting the broader landscape of genetic and epigenetic changes in putative tumor suppressors that could contribute to tumorigenesis. In this context, we applied for the first time Tuba-seq technology to systematically investigate the role of *Tbx2* genes and their effectors *in vivo*. Contrary to gene-centric models of tumor progression, we found that epigenetically dysregulated genes, such as the Tbx2 subfamily, can causally drive or slow tumor progression when deleted. This model provided highly precise and detailed quantification of tumor growth following the inactivation of our targeted genes in *Kras*-driven lung tumors.

In this study, genes associated with the TBX2 subfamily exhibited distinct roles at different stages of tumor progression. *Chd2* knockout inhibited both tumor progression and growth, while *Tnfaip3* knockout promoted tumor initiation and overall growth. Interestingly, cells deficient in *Tbx3*, *Tbx4*, *Tbx5*, and *Atf3* demonstrated minimal effects on tumorigenesis within our experimental model. Consistent with most studies, *Tbx2*-deficient cells exhibit suppressed growth capacity, yet its knockout significantly contributed to both tumor initiation and advanced progression—two aspects of TBX2’s role in tumorigenesis that remain understudied. Tbx2 is known to physically interact and suppress the transcriptional activity of *Egr1 – the most significant TSG identified in our study. Egr1* is a zinc-finger transcription factor with multifunctional roles in proliferation, stress responses, and apoptosis(22). In cancers such as rhabdomyosarcoma and breast cancer, TBX2 exploits this interaction to target several carcinogenic genes, including NDRG1, a protein involved in cell differentiation, apoptosis, and senescence (Fig. 1A) (23,24).

Beyond its relationship with TBX2, *EGR1* has been extensively studied in cancer biology. It is activated through the MAPK signaling pathway in response to various stimuli, such as growth factors, tumor necrosis factor, hypoxia, inflammatory signals, ionizing radiation, and reactive oxygen species. Once activated, *Egr1* can either promote or suppress the expression of its target genes, influencing transcriptional regulation. As a tumor suppressor in gliomas and melanocytomas, Egr1 upregulates p21Waf1/Cip1 to induce apoptosis(23). Additionally, *Egr1* enhances tumor cell death by directly upregulating tumor suppressors like NSAID-activated gene 1 (*NAG1*) and *PTEN*(*21*). In the context of immune regulation, *Egr1* has been the subject of conflicting reports, with its role oscillating between immunostimulatory and immunosuppressive effects depending on the cellular and environmental context. In inflammatory lung diseases, EGR1 is recognized as a master regulator of transcription, promoting the expression of genes critical for immune signaling and inflammatory cell activation(24,25). Conversely, within macrophages, EGR1 has been found to interact indirectly with the NuRD complex, leading to chromatin deacetylation and the suppression of inflammatory enhancer activation, thereby curbing excessive inflammatory responses(24). Despite its importance, EGR1 remains largely understudied in lung cancer, with existing research primarily limited to clinical surveys and in vitro experiments (26,27). For example, *EGR1* expression has been shown to predict *PTEN* levels and patient survival after surgical resection in NSCLC, with lower *EGR1* levels correlating with poorer outcomes(27). In this study, we present the first lung-specific *Egr1*-knockout mouse model, emphasizing its critical role at various stages of LUAD development, particularly in the context of Kras-driven tumorigenesis. *Egr1*-deficient cells demonstrated enhanced tumor initiation, growth, and progression, with effects exceeding those observed with the loss of *Rb1*, a key tumor suppressor in LUAD. Our transcriptomic analysis revealed that the tumorigenic effects of Egr1 deficiency are linked to an upregulation of immune-related genes and a molecular signature suggestive of T cell exhaustion in the tumor microenvironment. However, further investigation is needed to fully understand how *Egr1* deficiency in epithelial cells contributes to the onset or exacerbation of T cell exhaustion, offering new insights into the complex immune dynamics within tumors.

In conclusion, our study identifies *Egr1* as a key player in lung cancer progression and immune modulation, positioning it as a promising target for immunotherapy or a potential biomarker. We note that this discovery would not have been made were this study limited to (i) *in vitro* or (ii) clinical exome-level sequencing analyses, or (iii) a single gene within the TBX2 signaling cascade. Hence, this study underscores the indispensable value of systematic *in vivo* genetic models of cancer, which provide a more comprehensive understanding of gene function and mechanism.

## Methods

### 1 Mice and Tumor initiation

All animal experiments conducted in this study received approval from the Institutional Animal Care facility at Case Western Reserve University, under protocol number 2020-0099. Kras^LSL-G12D,Rosa26CAG-LSL-Cas9-GFP^ mice have been described(28,29). Mice were on a mixed BL6/129 background. Approximately equal numbers of males and females were used for each experiment. The corresponding figure legends specify the number of mice used in each experiment. Lung tumor initiation was achieved through intratracheal administration of viral-Cre vectors, following established protocol(30). Ketamine/xylazine drugs was used for anesthesia before the intubation step and the atipamezole was used directly afterwards as a reversal drug. The assessment of tumor burden encompassed various methodologies, including fluorescence microscopy, lung weight measurements, and histological analyses, all of which are comprehensively detailed in the appropriate sections of this study.

### 2 Design and generation of Lenti-sgRNA/Cre vectors

We designed lentiviral vectors to encompass Cre alongside single guide RNAs (sgRNAs) targeting specific genes under investigation (putative tumor suppressors and controls) in lung adenocarcinoma. These genes included *Tbx*, *Tbx3, Tbx4, Tbx5, Egr1, Tnfaip3, Chd2, and Atf3* (Supplementary Table. S1). Additionally, vectors carrying inert guides sg*Neo1* and sg*Neo2* were generated as internal controls. For validation purposes, sg*Rb1* served as a positive control, while an sgRNA targeting *Pcna*, an essential gene, acted as a negative control. The sgRNA sequences were meticulously crafted using CRISPRpick. We identified all feasible 20-bp sgRNAs with an NGG protospacer-adjacent motif (PAM) targeting each tumor-suppressor gene. These sgRNAs were evaluated for predicted on-target cutting efficiency through an sgRNA design/scoring algorithm. From this analysis, two unique sgRNAs per tumor-suppressor gene were selected, prioritizing those with the highest predicted cutting efficiencies. Preference was given to sgRNAs targeting exons conserved across all known splice isoforms (ENSEMBL), positioned closest to splice acceptor or donor sites, located earliest in the gene-coding region, and situated upstream of annotated functional domains (InterPro; UniProt) (Supplementary Table. S1).

The Lenti-U6-sgRNA-sgID-barcode-Pgk-Cre vector was cloned in collaboration with Genewiz. Initially, the sgRNA sequence of the pLenti-sgRb1/Cre vector (Addgene # #89647) was replaced by GCGAGGTATTACCGGCGTATCATCCGCG using site-directed mutagenesis, thereby creating pLenti-BaeI-Pgk-Cre. This replacement sequence incorporated a recognition site for the type IIS restriction endonuclease BaeI, facilitating swift replacement of the sgRNA sequence. Subsequently, forward and reverse single-stranded oligonucleotides containing the sgRNA sequence and complementary overhangs were annealed and ligated into the BaeI-linearized pLenti-BaeI-Pgk-Cre vector utilizing T4 DNA ligase. The barcode oligo primer, featuring the 8-nucleotide sgID sequence and 20-nucleotide degenerate barcode, was generated and ligated into the vectors according to established protocols(16).

### 3 Production, purification, and titering of lentivirus

To prevent barcode-sgRNA uncoupling due to lentiviral template switching during reverse transcription of the pseudo-diploid viral genome, individual barcoded Lenti-sgRNA/Cre vectors were generated separately(31). HEK293T cells were cultured in Dulbecco’s Modified Eagle Medium supplemented with 10% Fetal Bovine Serum and transfected with each of our barcoded Lenti-sgRNA/Cre plasmids, along with pCMV-VSV-G (Addgene #8454) envelope plasmid and pCMV-dR8.2 dvpr (Addgene #8455) packaging plasmid, using polyethylenimine. Following transfection, the cells were treated with 20 mM sodium butyrate after 8 hours, the culture medium was changed 24 hours later, and supernatants were collected at 36 and 48 hours post-transfection. Cell debris was removed using a 0.45 μm syringe filter unit (Millipore SLHP033RB), and each lentiviral vector was concentrated by ultracentrifugation (25,000 r.p.m for 1.5 hours at 4°C). The concentrated virus was resuspended in PBS and stored at −80°C. Vector titers were determined by transducing Rosa26LSL-YFP mouse embryonic fibroblasts with 10μl of each Lenti-sgRNA/Cre vector, assessing the percentage of YFP-positive cells via flow cytometry, and normalizing the titer to a lentiviral preparation with a known titer. At least 50k cell were sorted per sample. The cell line was confirmed negative for mycoplasma contamination. Lentiviral vectors were thawed and pooled immediately prior to administration to mice.

### 4 Genomic DNA Isolation from Mouse Lungs

Genomic DNA extraction was carried out from bulk tumor-bearing lung tissue obtained from each mouse. Prior to tissue homogenization, six benchmark control cell lines, comprising approximately 1×10^5 cells for three cell lines and 100 cells for three cell lines, each carrying unique sgID-BCs, were introduced (“spiked-in”) to every sample, as previously described (36). This addition facilitated the subsequent calculation of the absolute number of neoplastic cells in each tumor based on the sgID-BC reads. The entire lung tissues or right lobe from each mouse, along with the benchmark cell lines, underwent homogenization using a gentleMACS Dissociator. Homogenization was performed in 6 ml lysis buffer (100 mM NaCl, 20 mM Tris, 10 mM EDTA, 0.5% SDS) supplemented with 200 μl of 20 mg/ml Proteinase K (Life Technologies, AM2544). The homogenized tissue was incubated at 55 °C overnight. Subsequently, DNA extraction was carried out through phenol–chloroform extraction and ethanol precipitation from approximately 1/6th of the total lung lysate using standard protocols. DNA concentrations were determined using a nanodrop.

### 5 Library Construction and Sequencing of sgID-BC for Quantitative Analysis

Libraries were prepared through the amplification of the sgID-BC region from 32 μg of genomic DNA per mouse. The sgID-BC region of the integrated Lenti-sgRNA-BC/Cre vectors underwent PCR amplification using primer pairs that include TruSeq Illumina adapters and a 5′ multiplexing tag (TruSeq i7 index region indicated by Bold). This amplification utilized a universal forward primer primer (5′ AATGATACGGCGACCACCGAGATCTACACTCTTTCCCTACACGACGCTCTTCCGATCTGCGCACGTCTGCCGCGCTG 3′) and a unique reverse primer (5′ CAAGCAGAAGACGGCATACGAGAT**NNNNNN**GTGACTGGACTTCAGACGTGTGCTCTTCCG ATCCAGGTTCTTGCGAACCTCAT 3′).

A single-step PCR amplification of sgID-BC regions was employed, proving to be a highly reproducible and quantitative method for determining the neoplastic cell count in each tumor. For each mouse, eight 100 μl PCR reactions per sample (4μg DNA per reaction, 32μg per mouse) were conducted using Q5 High-Fidelity 2x Master Mix (New England Biolabs, M0494X). The resulting PCR products were purified with Agencourt AMPure XP beads (Beckman Coulter, A63881) using a double size selection protocol.

The concentration and quality of the purified libraries were assessed using the Agilent 2100 Bioanalyzer (Agilent Technologies, G2939BA). The libraries were pooled based on lung weight for even sequencing depth distribution, cleaned up, size-selected using AMPure XP beads, and sequenced on the Illumina® HiSeq 2500 platform, generating Paired-End 150 bp reads (PE150) (Novoegene). To enhance sequencing diversity and improve quality, 15-25% PhiX control DNA was added to the library at the beginning of the sequencing reads.

### 6 Quantification of tumor cell number from Tumor Barcode sequencing

High quality PE150 reads were each aligned to the known reference sequence of the lentiviral construct encapsulating 10 nucleotides flanking the dual barcodes on either side. Alignment to the reference sequence was performed on both the forward and reverse read using a Striped Smith Waterson algorithm to ensure lossless extraction of putative barcodes from every read(32). A prespecified scoring matrix of: (i) nucleotide match = +4, (ii) nucleotide mismatch = −2, (iii) gap open = −6, (iv) gap extension = −1 was used. Barcodes were then extracted from each the forward and reverse read alignments with mismatching bases annotated as ‘N’. Barcodes were then ‘dereplicated’ (tallied) and barcode tallies were removed if (1) the forward and reverse barcode lengths did not match, or (2) the mean alignment score of the entire pileup was below 50%. This lossless dereplication of barcodes followed by filtering based on the global properties of the barcode pileup minimized biases that sequencing fidelity imparts on pileup quantity (e.g. barcodes with homopolymer runs generally exhibit lower PHRED scores), while still ensuring that only true lentiviral barcodes are tallied.

The rate of recurrent read errors was estimated and used to remove spurious barcodes from pileups using an eight parameter error model: six symmetric substitution rates to/from A, T, C, G, N; and an Insertion and Deletion rate. These parameters were estimated once from the median rate of observed recurrent read errors descending from the 100 largest barcode pileups in every mouse in this study. The probability that every barcode in this study was a recurrent read error was then estimated by calculating a rate statistic <u>λ</u> from each barcode’s nearest neighbor barcode size multiplied by the estimated error rate for the difference between the two barcodes in question (e.g. substitution or InDel). A Poisson distribution was then used to estimate the likelihood that a pileup was a recurrent read error based on this rate statistic and pileups with a likelihood >10^-10^ were filtered from this study (Supplementary Fig. S1B, C and D). This filtering probability has been used previously(16,33).

GC bias during PCR amplification and the likelihood of ‘barcode collisions’ were then estimated for every mouse and Lenti-sgRNA/mBC pool; however, minimal variation across the experiment was observed. A ‘barcode collision’ rate constitutes the likelihood that two lentiviruses with the exact same barcode pair transduce different cells in the same mouse. The probability of barcode collisions can be estimated based on the frequency with which the same barcodes appear in separate mice in the study (high diversity barcodes appear only once in the entire study, whereas lower diversity barcodes may be observed in two or three mice) The probability of barcode collisions can bias growth estimates if Lenti-sgRNA pools are barcoded with varying efficiency, however in this study we did not observe appreciable differences in barcode diversity across sgRNAs. PCR amplification bias was also not observed, as seen previously(16).

Absolute tumor numbers were then estimated for every non-filtered barcode pileup in every mouse by simply dividing read number by the median read number observed for the three spike-in barcodes and multiplying by the known cell number of these barcodes (1 × 10^5^). A minimum cell cutoff of 400 cells yielded tumor size profiles that were highly-reproducible across the technical replicates in our study, and minimized variation in median tumor size between the two inert sgRNAs (Neo1 and Neo3).

### 7 Comprehensive statistical assessment of tumor progression using TuBa-seq

Previous TuBa-seq analyses found that a single summary statistic cannot explain differences in tumor progression observed between different tumor genotypes. In particular, both the *quantity* of transduced lineages and size spectrum of distinct genotypes reproducibly varies (e.g. Stk11 loss dramatically increases mean tumor size without appreciably altering tumor number, while Pten loss dramatically increases tumor number with only a moderate increase in mean tumor size(34). Size spectrums then reproducibly differ in both the central tendency (mean/median) of growth, as well as the preponderance of exceptionally-large tumors [NatMethods], which cannot be explained by any single random process. Because malignancies are only observed at later time points (and most transduced lineages remain premalignant), it is imperative to quantify both the immediate effects of gene loss (mean and/or 50^th^ – 95^th^ percentile) and the ability of a genotype to potentiate clinically-relevant disease (‘heavy-tail’ tumor burden and/or 99^th^ – 99.99^th^ percentile).

Total tumor quantity, percentiles of tumor size from the 10^th^ to 99.99^th^ percentiles, and parametric summary statistics of size profiles were all calculated for every sgRNA pool in every mouse as described previously(16). Central tendency was estimated using the MLE of mean for a Log-Normal distribution, while deviation from a Log-Normal distribution in the large ‘heavy’ tail of tumor sizes using(35). The minimum size cutoff for the Pareto tail of tumor burden was calculated relative to median tumor size of sg*Inerts* and inferred to be 44.73 times the median for this study. All statistics for active sgRNAs were then divided by their respective statistic for sg*Inerts* within each mouse, and then averaged across all mice in a particular experimental cohort. P-values and confidence intervals for percentiles and parametric statistics were then calculated using two million iterations of the Bootstrap resampling method. P-values for tumor number (tumor initiation) were calculated using Fischer’s Exact Test.

### 8 Total RNA Isolation Protocol from Mice Lung tissues and RNAseq

Tumor-bearing lung lobes from mice intubated with Lenti-sg*Egr1*/Cre or Lenti-sg*Inert*/Cre pools were mixed with TRIzol reagent (Invitrogen Life Technologies) and homogenized using a gentleMACS Dissociator (Miltenyi Biotec). Subsequently, 1 ml of the lysate was treated with 0.2 ml of chloroform per 1 ml of TRIzol and vigorously shaken for 15-30 seconds, followed by incubation at room temperature for 2-3 minutes to induce phase separation. The mixture was then centrifuged at 12,000-16,000 x g for 15 minutes at 4°C. The RNA-containing aqueous phase was carefully transferred to a new tube, and RNA purification was performed using the Qiagen RNeasy Mini Kit (Qiagen #74104) according to the manufacturer’s protocol. RNA quality was assessed based on RNA integrity numbers (RINs) using a 2100 Bioanalyzer. Sequencing was performed using NovaSeq X PlusSeries (PE150) reading 12 G raw data per sample.

### 9 Histology and Immunohistochemistry

Lung tissue samples were obtained from 1-3 mice per cohort, transduced with the Lenti-sgTSPool/Cre pool or individual sgRNAs (sg*Egr1* and sg*Inerts* pools). The collected lobes were fixed in 4% paraformaldehyde for 12 hours, followed by storage in 1X PBS at 4°C. Subsequently, tissues were paraffin-embedded and sectioned at the University Hospital Histology Core Facility into 4-micrometer sections. Hematoxylin and eosin (H&E) staining was conducted for visualization, and tumor sizes were assessed by measuring the longest diameter of each tumor in the H&E stained sections using ImageJ software. Immunohistochemical staining was performed using specific antibodies: anti-TTF-1/Nkx2–1 (1:200, abcam, AB76013) anti-CD3 (1:750, AB135372), anti-CD4 (1:1000, AB183685), antiCD8 (1:1000, ab217344), anti-CD19 (1:1000, ab245235), anti-F4/80 (1:100, Invitrogen# 14-4801-85) and anti-Egr1 (1:200, ab6054). Statistical analysis was performed using ImageJ software, with p-values calculated using the Mann-Whitney U test.

### 10 RNAseq analysis

Raw sequence reads were processed to remove contaminant DNA, PCR duplicates, and adaptor sequences. Reads with a quality score below Q20 were excluded using CLC Genomics Workbench (v22.0.2). Paired-end reads were then assembled and aligned to the latest human genome reference (Homo sapiens, GRCh38). Transcript assembly and abundance quantification were conducted via StringTie(36). After FPKM Normalization with Ballgown(37), different gene expression analysis was conducted with edgeR’s exact test (edgeR v4.2.1)(38). Pairwise comparisons were performed with edgeR’s decideTestsDGE function and Benjamini-Hochberg (BH)-adjusted p-value cutoff of 0.05 was used to obtain differentially expressed genes (absolute log2[Fold-Change] ≥ 1). Gene set enrichment analysis (GSEA) was performed using the murine hallmark gene set from MSigDB (v7.5.1)(39); significantly enriched pathways had BH-adjusted p-value ≤ 0.05 and minimum gene set size of 10. Data visualizations were conducted through R (v4.4.1) packages. Significantly upregulated genes underwent Gene Ontology (GO) enrichment analysis using the Metascape tool. Statistically enriched GO terms were identified using cumulative hypergeometric p-values and enrichment scores. Then, a network-based approach was used to identify the most central genes in immunity-related pathways. Specifically, differentially upregulated genes from the top 18 significantly enriched and immunity-related pathway clusters were analyzed for Maximal Clique Centrality (MCC) using the CytoHubba plug-in of Cytoscape. A comprehensive set of T-cell exhaustion markers was curated from multiple resources, including CellMarker 2.0 and 10X Genomics. Differential expression of 55 curated markers was assessed between Egr1-KO and Control samples, and their expression profiles were visualized in a heatmap.

### 11 *Egr1* Knockout and Expression Analysis Using Publicly Available DepMap and PCAWG Datasets

The impact of EGR1 knockout was analyzed using RStudio with data from DepMap CRISPRGeneEffect dataset, which was integrated with the model dataset to classify cancer types, and with the OmicsSomaticMutations dataset to identify cell lines harboring KRAS mutations. To determine the significance of EGR1 knockout effects, we applied cutoffs of −0.5 and 0.5, as suggested by DepMap. *EGR1* expression in clinical samples was evaluated using the Pan-Cancer Analysis of Whole Genomes (PCAWG) dataset. Expression levels in normal lung tissue were compared to those in lung adenocarcinoma by filtering samples based on histology and using DCC project codes to identify normal tissue. Statistical significance between the two groups was calculated using the Wilcoxon signed-rank test.

## Author Contribution

AK and CM conceptualized the study, designed the experiments, and supervised the project. AK performed the experiments. TD, MP, RO and CM analyzed the data. ZF, ZW, JM and BG assisted with data analysis and interpretation. AK, MR, XW, MB, CD, TD, MP and YW contributed to the development of the methodology and provided critical resources. AK wrote the manuscript. CD, BG, JS, TF, RO and ZW provided critical input and edits to the manuscript. All authors reviewed and approved the final version of the manuscript.

## Acknowledgment

This work was supported by grants from the National Institutes of Health (No. R01CA271540, R00CA226506 to CM; No. R01CA196643, R01CA264320, R01CA260629, P50CA150964, and P30 CA043703 to ZW

## Supplementary figures’ legends

Supplementary Figure1: (A) Pearson correlation coefficient (r) and two-tailed P-value indicate a correlation between neoplastic cell count (a measure of tumor burden) and lung weight. Each dot represents an individual mouse. (B) Total cell burden for KC mice after 6 and 20 weeks of tumor growth. Total cell number for each mouse calculated by summing absolute cell # of every tumor within a mouse assigned to an Inert or Tumor Suppressor Gene (TSG) knockout guide RNA. The mean burden for each mouse and 95% CI estimate of the mean, determined by bootstrap resampling, are denoted by horizontal and vertical bars, respectively. In general, TSG knockout sgRNAs increase total tumor burden, consistent with their purported carcinogenic role. (C) Inferred error rates of TuBa-seq analysis pipeline. Substitutions and single-base InDel (Insertion or Deletion) mutations can lead to spurious tumor calls. Hence, we infer the rate of each error by measuring the median number of reads belonging to barcode pileups within one error of the top 100 barcodes observed in every mouse. For example, a large pileup of 100,000 ACGT barcodes might spawn recurrent read errors with a C → G substitution such that multiple A<u>G</u>GT barcodes are observed. These inferred error rates are then used to eliminate spurious tumor calls (Methods). The rate of substitutions which lead to a consistent barcode error in both the forward and reverse sequencing of the Paired-end read arise at a rate of approximately 10^-5^ nucleotide^-1^. This error rate – and profile favoring C → G substitutions, is consistent with the expected error rate for PCR barcode amplification by Hi-Fideldity Q5 DNA polymerase (Methods). (D) Sequencing error rates are inferred by calculating the rate at which the forward read sequence of a barcode mismatches the reverse read sequence. Sequencing error rates are somewhat consistent with mean PHRED scores, albeit somewhat greater than expected rates. (E) Inferred InDel error rates of all barcodes in this study are much lower than substitution and sequencing error rates. (F) Mean effect of gene knockout of each sgRNA in KC mice at 20-weeks post tumor initiation. LN Mean of each gene for both of the high-specificity sgRNAs detailed in Table S1. Growth effects of the two guide RNAs are similar, but less consistent than at six weeks. This may be because size distributions at 20 weeks are heavy-tailed and therefore poorly explained by summary statistics for a Log-Normal distribution.

Supplementary Figure2: (A)Estimates of mean tumor size, assuming a lognormal tumor size distribution, identified sgRNAs that significantly increase growth in KC mice. Bonferroni-corrected, bootstrapped p-values are shown. p-values < 0.05 and their corresponding means are bold. (B) Analysis of relative tumor sizes in KRAS driven tumors after 6 and 20 weeks of intubation. Relative size of tumors at the indicated percentiles is unmerged data of individual sgRNA from each cohort of mice, normalized to the average size of sgInert tumors. Percentiles significantly different from sgInert are in color. The darker the shade of color the larger the percentile, as shown in the legend in grayscale. Error bars denote 95% confidence intervals determined by bootstrap sampling.

Supplementary Figure3: (A) Tumor size in *Egr1*-deficient lung tumors was compared to controls by measuring the longest diameter of each tumor. The number of tumors analyzed is shown on the X-axis. P-values were calculated using a one-tailed Mann-Whitney U test, with horizontal lines representing the median and whiskers indicating the minimum and maximum values. (B) Principal Component Analysis (PCA) plot of transcriptome of tumor-bearing *Egr1*-knockout lungs and control mice demonstrating clear separation between the two groups. (C) Gene Set Enrichment Analysis on DSigDB using clusterProfiler between *Egr1*-deficient lung and control samples. Top 3 most enriched pathways in EGR1-KO group are Allograft Rejection, Interferon Alpha, Interferon Gamma. (D) *EGR1* expression in clinical samples was evaluated using the Pan-Cancer Analysis of Whole Genomes (PCAWG) dataset. Expression levels in normal lung tissue were compared to those in lung adenocarcinoma by filtering samples based on histology and using DCC project codes to identify normal tissue. Statistical significance between the two groups was calculated using the Wilcoxon signed-rank test.

Supplementary Table S1: Lenti-sg*RNA* vectors used for gene targeting. The table lists the sgRNA sequences used to target each of the genes analyzed in this study and their corresponding sgIDs. sgRNAs targeting *Rb1* (Addgene #89647), *Neo1* (Addgene #67594), *Neo3* (Addgene #89653), and *Pcna*, has been adopted from previous studies(16).

Supplementary Table S2: Lung cancer cell lines harboring KRAS mutations adopted from OmicsSomaticMutations dataset.

