## Supplementary figures and images for "*In Vivo* Multiplexed Modeling Reveals Diverse Roles of the TBX2 Subfamily and *Egr1* in *Ras*-Driven Lung Adenocarcinoma"

### Supplemental figure 1

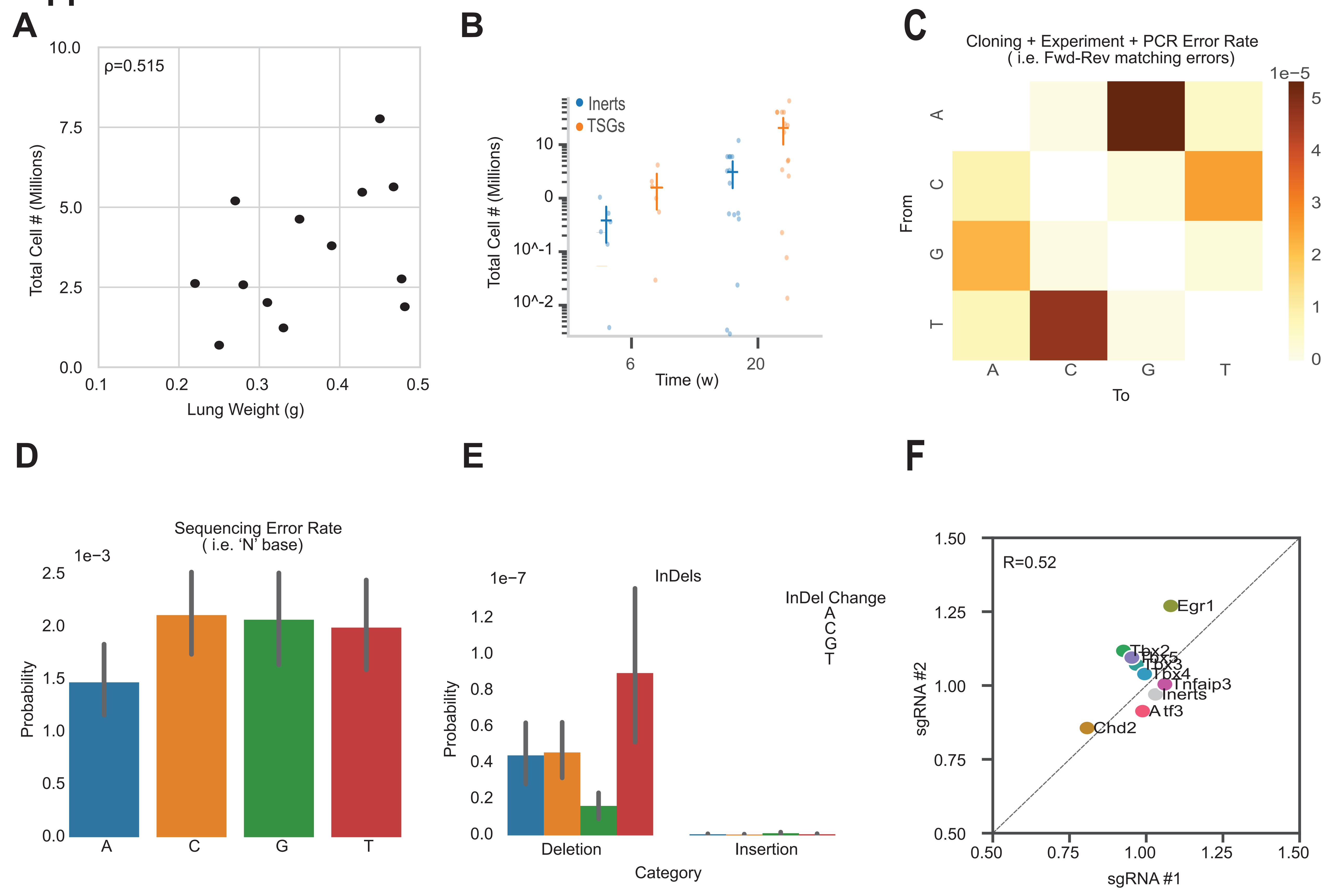

### Supplemental figure 3

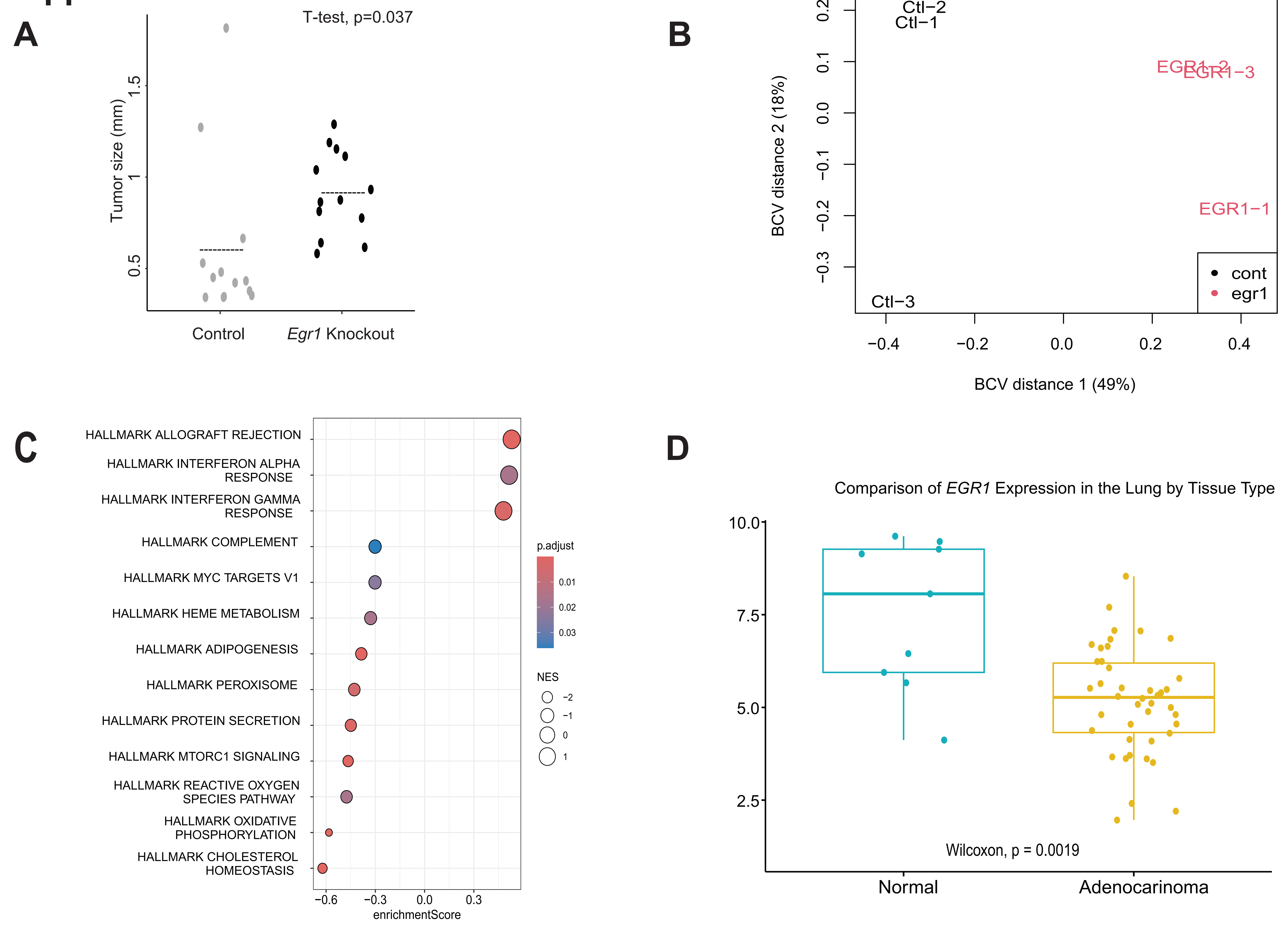
