## Supplemental figure 2 for "*In Vivo* Multiplexed Modeling Reveals Diverse Roles of the TBX2 Subfamily and *Egr1* in *Ras*-Driven Lung Adenocarcinoma"

**Supp-2**

**A**

| sgRNA | 6 weeks: Mean (relative to inerts) | 6 weeks: P-Value | 20 weeks: Mean (relative to inerts) | 20 weeks: P-Value |
|---|---|---|---|---|
| *Atf3* | 0.885 | 0.164 | 0.948 | 0.274 |
| *Chd2* | 0.789 | 0.009 | 0.832 | 0 |
| *Egr1* | 1.340 | 0 | 1.129 | 0.004 |
| *Tbx2* | 0.707 | 0 | 0.999 | 0.983 |
| *Tbx3* | 0.933 | 0.403 | 1.026 | 0.602 |
| *Tbx4* | 1.038 | 0.662 | 1.024 | 0.603 |
| *Tbx5* | 0.903 | 0.235 | 1.019 | 0.710 |
| *Tnfaip3* | 1.188 | 0.028 | 1.035 | 0.472 |
| *Rb1* | 1.115 | 0.166 | 2.501 | 0 |
| *Pcna* | 0.863 | 0.098 | 0.940 | 0.198 |

**B**

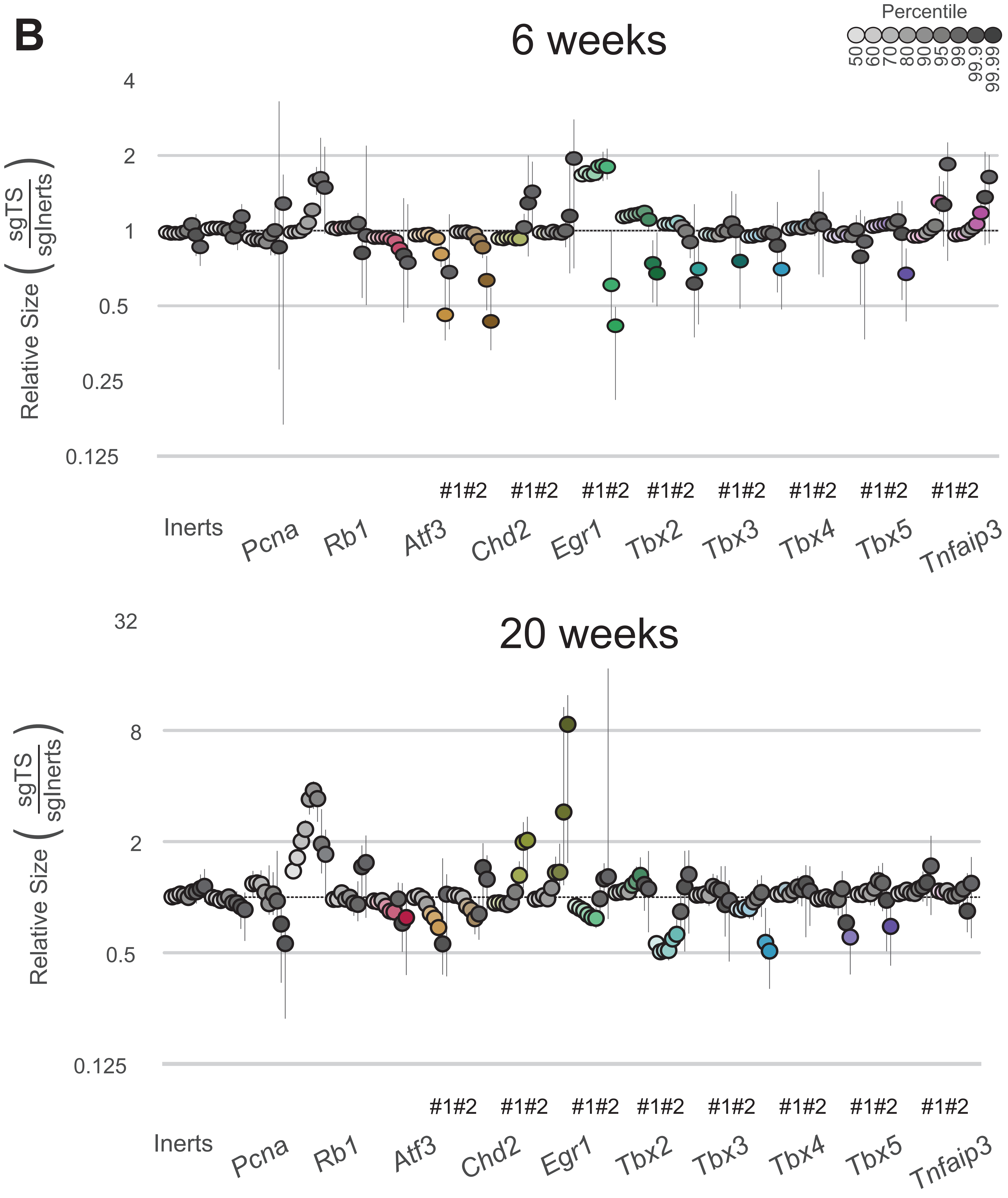
