## Supplemental Table-1 for "*In Vivo* Multiplexed Modeling Reveals Diverse Roles of the TBX2 Subfamily and *Egr1* in *Ras*-Driven Lung Adenocarcinoma"

| sgRNA target | sgRNA number | sgRNA Sequence | PAM Sequence |
|---|---|---|---|
| *Tbx2* | #1 | GCTCGCACGATGTGGAATCG | CGG |
| *Tbx2* | #2 | GTCCGGCCACAGGGGAACAG | TGG |
| *Tbx3* | #1 | GAGCACCTCACTTTAAACGG | AGG |
| *Tbx3* | #2 | CATCATGGATCAGTTAGTGG | GGG |
| *Tbx4* | #1 | CTTGTAGCGATGGTCATCTG | CGG |
| *Tbx4* | #2 | CCCGGATTCTCCTGCCACCG | GGG |
| *Tbx5* | #1 | TGGCTGAAGTTCCACGAAGT | GGG |
| *Tbx5* | #2 | CGAAACCTGAGAGTGCTCTG | GGG |
| *Egr1* | #1 | GAGGATTGGTCATGCTCACG | AGG |
| *Egr1* | #2 | GTTATCCCAGCCAAACGACT | CGG |
| *Tnfaip3* | #1 | ACTGACAAGCTGCATGCATG | AGG |
| *Tnfaip3* | #2 | AACCATGCACCGATACACGC | TGG |
| *Chd2* | #1 | AAGCAACCTAAGATTCAGCG | TGG |
| *Chd2* | #2 | GTCTTATATTCACAGCACAT | AGG |
| *Atf3* | #1 | TCAAATACCAGTGACCCAGG | AGG |
| *Atf3* | #2 | GGCGGTCGCACTGACTTCTG | AGG |

| Exon Number | Target Cut Length | Target Cut % | On-Target Efficacy Score | sgID |
|---|---|---|---|---|
| 3 | 699 | 32.7 | 0.6592 | TTGGCAAC |
| 2 | 591 | 27.7 | 0.6211 | AATGCGTG |
| 2 | 405 | 18.7 | 0.6961 | TTTACCCG |
| 1 | 205 | 9.5 | 0.6828 | TTGCCTGT |
| 3 | 382 | 23 | 0.6082 | TGAGCTTG |
| 4 | 484 | 29.2 | 0.5905 | TTGTCCGA |
| 3 | 206 | 13.2 | 0.6608 | CAGTCGTA |
| 2 | 93 | 6 | 0.7005 | ATCGTTGC |
| 2 | 476 | 29.7 | 0.6483 | CGATTAGG |
| 2 | 366 | 22.8 | 0.6527 | CTTACGGT |
| 3 | 314 | 13.5 | 0.6959 | TTCCTCCT |
| 2 | 139 | 6 | 0.5872 | GCGGAATA |
| 7 | 593 | 10.8 | 0.684 | TTCCAAGC |
| 4 | 315 | 5.7 | 0.4723 | TCAGTTCG |
| 2 | 78 | 14.3 | 0.7489 | ACTTGGTC |
| 2 | 37 | 6.8 | 0.6238 | TAGCCTGT |
