## Supplemental Table-2 for "*In Vivo* Multiplexed Modeling Reveals Diverse Roles of the TBX2 Subfamily and *Egr1* in *Ras*-Driven Lung Adenocarcinoma"

|  | KRAS Lung cancer Cell line name |
|---|---|
| 1 | NCI-H2887 |
| 2 | Calu-6 |
| 3 | NCI-H2122 |
| 4 | HCC-461 |
| 5 | NCI-H647 |
| 6 | NCI-H1944 |
| 7 | Lu-65 |
| 8 | LU99 |
| 9 | NCI-H460 |
| 10 | NCI-H1792 |
| 11 | Calu-1 |
| 12 | NCI-H2030 |
| 13 | LCLC-97TM1 |
| 14 | NCI-H441 |
| 15 | COR-L23 |
| 16 | NCI-H1355 |
| 17 | HCC-44 |
| 18 | IA-LM |
| 19 | SW 1573 |
| 20 | A549 |
| 21 | NCI-H2291 |
| 22 | A427 |
| 23 | RERF-LC-Ad2 |
| 24 | NCI-H727 |
| 25 | SHP-77 |
| 26 | RERF-LC-Ad1 |
| 27 | NCI-H1373 |
| 28 | MOR/CPR |
| 29 | NCI-H358 |
| 30 | HOP-62 |
| 31 | HCC515 |
| 32 | NCI-H2009 |
| 33 | NCI-H23 |
| 34 | NCI-H1573 |
| 35 | NCI-H157-DM |
| 36 | NCI-H650 |
| 37 | NCI-H1155 |
| 38 | A549_CRAF_KD |
